# Efficient High-Throughput DNA Breathing Features Generation Using Jax-EPBD

**DOI:** 10.1101/2024.12.06.627191

**Authors:** Toki Tahmid Inan, Anowarul Kabir, Kim Rasmussen, Amarda Shehu, Anny Usheva, Alan Bishop, Boian Alexandrov, Manish Bhattarai

## Abstract

DNA breathing dynamics—transient base-pair opening and closing due to thermal fluctuations—are vital for processes like transcription, replication, and repair. Traditional models, such as the Extended Peyrard-Bishop-Dauxois (EPBD), provide insights into these dynamics but are computationally limited for long sequences. We present *JAX-EPBD*, a high-throughput Langevin molecular dynamics framework leveraging `JAX` for GPU-accelerated simulations, achieving up to 30x speedup and superior scalability compared to the original C-based EPBD implementation. *JAX-EPBD* efficiently captures time-dependent behaviors, including bubble lifetimes and base flipping kinetics, enabling genome-scale analyses. Applying it to transcription factor (TF) binding affinity prediction using SELEX datasets, we observed consistent improvements in *R*^2^ values when incorporating breathing features with sequence data. Validating on the 77-bp AAV P5 promoter, *JAX-EPBD* revealed sequence-specific differences in bubble dynamics correlating with transcriptional activity. These findings establish *JAX-EPBD* as a powerful and scalable tool for understanding DNA breathing dynamics and their role in gene regulation and transcription factor binding.

## Introduction

DNA breathing dynamics, characterized by the transient opening and closing of base pairs due to thermal fluctuations, play a crucial role in fundamental biological processes such as transcription initiation, replication, and DNA repair (Watson et al. 2013). These local strand separations facilitate access to genetic information and are influenced by thermal energy that disrupts the weak hydrogen bonds between complementary bases (Guéron, Kochoyan, and Leroy 1987).

To quantitatively study DNA breathing, theoretical models like the Extended Peyrard-Bishop-Dauxois (EPBD) model have been developed (Alexandrov et al. 2009; Peyrard and Bishop 1989). The EPBD model extends the original PBD model by incorporating sequence-specific stacking potentials, allowing for the analysis of sequence-dependent effects on DNA dynamics. This model provides single-nucleotide resolution and captures the nonlinear, highly cooperative nature of DNA breathing, enabling the detection of effects from even single base pair changes (Alexandrov et al. 2009). Traditional thermodynamic models, while useful for predicting melting temperatures, struggle to account for deviations in melting behavior observed in homogeneous and periodic DNA sequences (SantaLucia Jr 1998; Ares et al. 2005). Dynamic models like EPBD offer advantages by capturing long-range effects and providing insights into the initial stages of DNA melting, which are relevant for processes like protein binding and transcription initiation (Peyrard 2004).

Previously, we used Markov Chain Monte Carlo (MCMC) simulations in the `pyDNA-EPBD` framework to examine DNA breathing dynamics (**?**). While effective for sampling equilibrium properties, MCMC methods do not provide temporal information about the dynamics, such as bubble life-times and the kinetics of base pair opening (Frenkel and Smit 2002). To address this limitation, we have transitioned to Langevin molecular dynamics (LMD) simulations using the `JAX` library (Bradbury et al. 2018). LMD incorporates both deterministic forces from the potential energy landscape and stochastic forces representing thermal fluctuations (Allen and Tildesley 1989), allowing us to capture time-dependent behavior and study kinetic properties of DNA breathing. Leveraging `JAX`’s just-in-time compilation and GPU acceleration, we implemented a highly scalable LMD simulation framework optimized for performance. This advancement enables extensive simulations of longer DNA sequences and the capture of rare breathing events with high temporal resolution.

In this work, we detail our LMD simulation framework using `JAX`, compare its performance and advantages over the MCMC approach, and discuss the implications of our findings for understanding DNA breathing dynamics and their role in biological processes.

### The Langevin-EPBD Model

The EPBD model is a dynamic, highly nonlinear system that describes the transverse opening motions of the complementary strands of double-stranded DNA, known as DNA breathing dynamics (Peyrard and Bishop 1989). This model allows for the existence of breathing solutions—transient but relatively long-lasting openings of the DNA double helix—which are closely linked to the local bending tendency of the DNA. The trajectories derived from the EPBD model provide detailed information about the lifetimes of these transient DNA openings, known as bubbles—a level of detail not accessible through purely thermodynamic calculations (Alexandrov et al. 2006). By explicitly considering solvent conditions such as salt concentration, temperature, and DNA twist, the EPBD model can reveal bubbles with extended lifetimes (Alexandrov et al. 2010a,b). Another key advantage of the EPBD model is its resolution at the singlenucleotide level. In contrast, thermodynamic models typically require averaging over windows of 100–500 base pairs to calculate property profiles, which can obscure differences between closely related sequences. The EPBD model avoids this averaging, enabling the detection of effects from even single base pair changes.

The EPBD model is a quasi-two-dimensional nonlinear framework designed to describe the transverse opening motion of the complementary strands of double-stranded DNA. It accounts for the distinction between the two sides (right *u*_*n*_ and left *v*_*n*_) of the DNA double strand, offering a more nuanced representation of its dynamics. The Hamiltonian potential surface of the EPBD model is characterized by the summation of two key energy terms: the Morse potential and the stacking energy of the two neighboring base pairs (bps) at every bp of the input DNA fragment, see equation (1).

Mathematically, the total potential energy is expressed as:

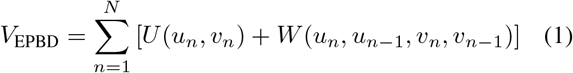

The Morse potential *U* (*u*_*n*_, *v*_*n*_) incorporates the hydrogen bonds between two bases belonging to opposite strands of the DNA. It also accounts for the repulsive interactions of the phosphate groups and the effects of the surrounding solvent. The parameters *D*_*n*_ and *a*_*n*_ in the Morse potential are specific to the *n*-th base pair, reflecting whether it is an A–T or G–C pair. This potential is given by:

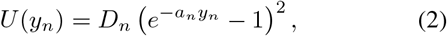

where:

- *u*_*n*_ and *v*_*n*_ are the transverse displacements of the complementary bases in the *n*-th base pair.
- 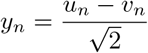 is the relative distance between the bases.
- *D*_*n*_ is the bond dissociation energy, and *a*_*n*_ determines the sharpness of the potential well.

This term models the balance between attractive forces (hydrogen bonding) and repulsive forces (phosphate interactions), capturing the effect of thermal fluctuations.

The **stacking potential** *W* (*u*_*n*_, *u*_*n*− 1_, *v*_*n*_, *v*_*n* −1_) describes the interaction between neighboring base pairs. It depends on the status variables of adjacent base pairs and accounts for the mechanical coupling between them. This term is expressed as:

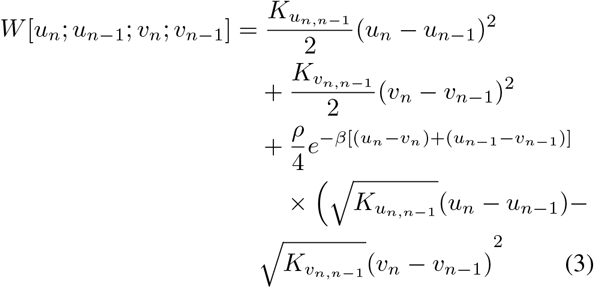

where:

- *u*_*n*_ and *u*_*n™* 1_ are the positions of the *n*-th and (*n™* 1)-th bases along one strand.
- *v*_*n*_ and *v*_*n™* 1_ are the corresponding positions on the complementary strand.
- 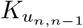 and 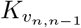 are the coupling constants determining the stiffness of the interactions between neighboring bases.
- *ρ* and *β* are parameters related to the strength and range of the interactions.

This term plays a critical role in maintaining the structural stability of the DNA double helix by ensuring cooperative behavior between adjacent base pairs.

In the EPBD model, the thermal dynamics of the *n*-th base pair are obtained through the Langevin equation:

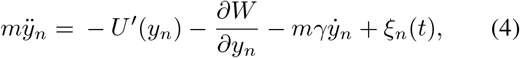

where:

- *m* is the mass of the base pair.
- *U* ^*′*^(*y*_*n*_) is the derivative of the Morse potential with respect to *y*_*n*_.
- 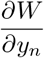 represents the derivative of the stacking potential with respect to *y*_*n*_, including contributions from neighboring base pairs.
- *γ* is the friction coefficient.
- *ξ*_*n*_(*t*) is a random force representing thermal fluctuations, modeled as white noise sampled from a standard Gaussian distribution.

We simulate the dynamics of double-stranded DNA at *T* = 310 K by numerically integrating the stochastic differential equation (4) with periodic boundary conditions using the Langevin dynamics method.

### Langevin dynamics

Langevin dynamics is a method used in molecular dynamics simulations to model the interaction between a system of interest and its surrounding environment, such as a solvent or heat bath. It introduces stochastic and frictional forces into Newton’s equations of motion to account for thermal fluctuations and energy dissipation. This method is particularly effective for studying systems at finite temperatures and exploring thermodynamic properties.

The general Langevin equation describes the Brownian motion of a particle:

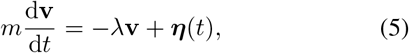

where:

- **v** is the velocity of the particle.
- *λ* is the damping coefficient, representing the frictional force due to the surrounding medium.
- *m* is the mass of the particle.
- ***η***(*t*) is a stochastic force (random noise) representing collisions with fluid molecules.

The random force ***η***(*t*) satisfies the statistical properties:

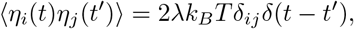

where:

- *k*_*B*_ is the Boltzmann constant.
- *T* is the temperature of the system.
- *δ*_*ij*_ is the Kronecker delta.
- *δ*(*t− t*^*′*^) is the Dirac delta function, indicating that the force at time *t* is uncorrelated with the force at any other time.

In molecular dynamics, the Langevin equation is adapted to describe the motion of particles in a system:

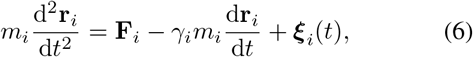

where:

- **F**_*i*_ is the deterministic force acting on particle *i*, derived from the system’s potential energy.
- *γ*_*i*_ is the friction coefficient for particle *i*, representing energy dissipation into the environment.
- ***ξ***_*i*_(*t*) is the stochastic force, with the same Gaussian properties as in the original Langevin equation.

Relating this to our EPBD model by comparing equations (4) and (6), we identify:

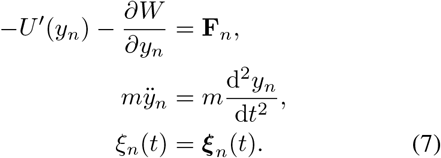

The Langevin equation allows the simulation of time-dependent behavior, showing how the DNA base pairs evolve under the combined effects of the EPBD potential, thermal fluctuations, and damping. The simulation involves solving the second-order ordinary differential equation (4). We do this using a numerical method called the second-order Runge-Kutta (RK2) (Ixaru et al. 2004) method. This numerical approach approximates the time evolution of base pair positions under the influence of the forces described by the EPBD model and the stochastic and damping terms from the Langevin equation.

To calculate the average displacement or opening profile for a given DNA sequence at a specific temperature, we utilize the EPBD model summarized in Algorithm 1 and illustrated in Figure 2. This involves solving for the relative distance *y*_*n*_ for each base pair. Subsequently, each base’s displacements *y*_*n*_ at selected time intervals are recorded. For each DNA sequence, we run at least 500 independent simulations to derive the average displacement/opening profile. Each simulation trajectory involves starting from a stationary position and numerically integrating equation (4) for a specified number of steps.

### High-Throughput Langevin EPBD

#### Algorithm 1: RK2 Method for Solving the PBD Model Equations of Motion

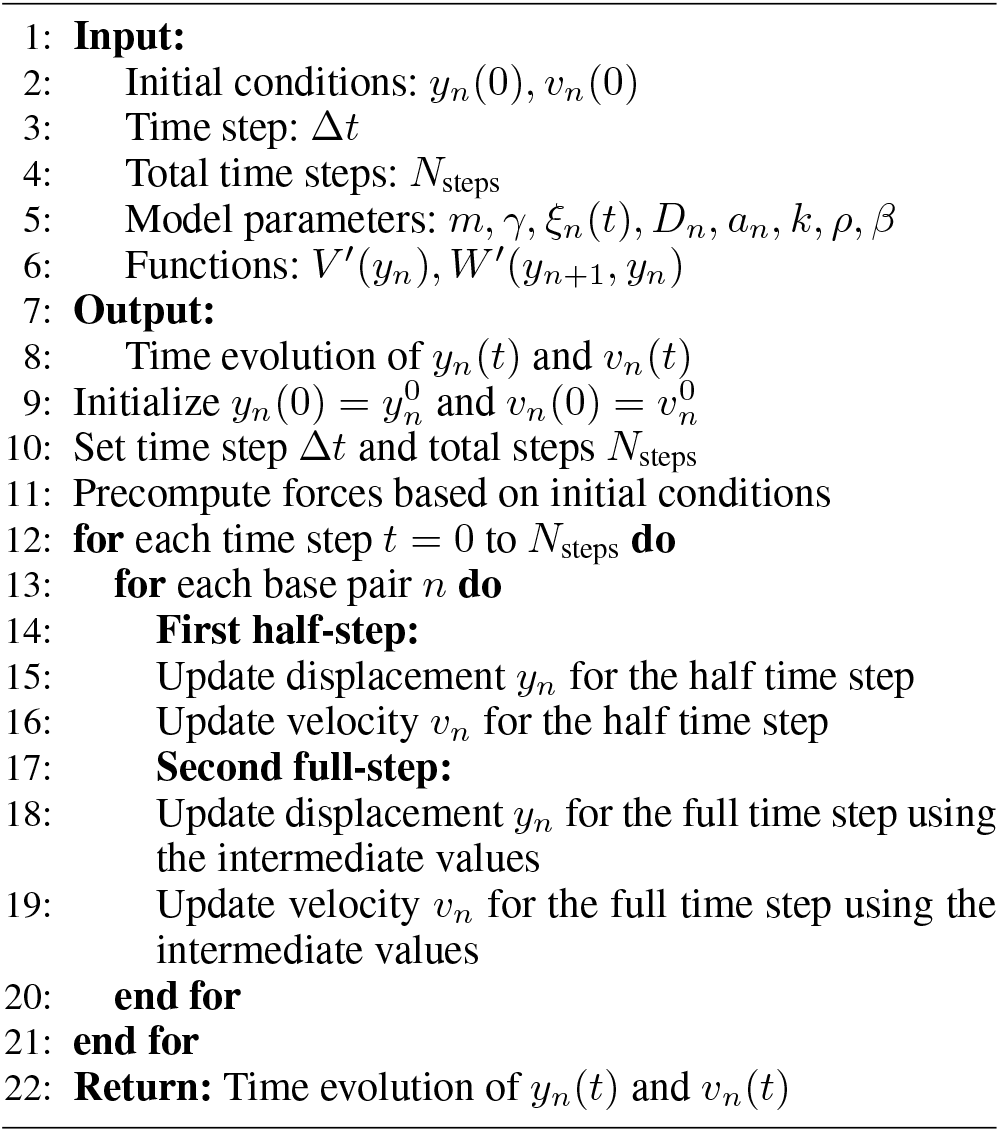

Numerical integration methods, such as those used in Langevin dynamics, require more computational steps and necessitate more independent simulations than MCMC methods to achieve the desired outputs. Following the implementation described in (Alexandrov et al. 2006), we employed `JAX` (Bradbury et al. 2018) as our implementation framework, in contrast to the *C* implementation used in (Alexandrov et al. 2006) or the native python implementation in (Kabir et al. 2023)

Although certain operations can be computationally expensive in `JAX`, the Accelerated Linear Algebra (XLA) framework significantly enhances computational efficiency by serving as a performance-optimizing compiler. XLA transforms high-level mathematical expressions into optimized kernels tailored for diverse hardware platforms, including GPUs. Additionally, `JAX` provides optimized computations for GPU programming models. Thus, vectorized code written in `JAX` can execute faster than code in other programming frameworks that operate solely on CPUs.

We refer to our Langevin-EPBD implementation using `JAX` as *JAX-EPBD*. The strength of *JAX-EPBD* lies in its unparalleled ability to process multiple sequences concurrently, a capability critical for efficient Langevin simulations on GPUs. Unlike other popular frameworks such as PyTorch or TensorFlow, which typically support only a single instance of simulation on GPUs at a time due to constraints in their design, `JAX` uniquely enables concurrent simulation instances. This is achieved through its highly optimized `vmap` (vectorized mapping) and `pmap` (parallel mapping) functions, which allow for seamless vectorization and parallel execution across GPU cores. By leveraging `vmap`, *JAX-EPBD* processes batches of sequences simultaneously, ensuring that each sequence is treated independently while sharing GPU resources efficiently. Furthermore, `pmap` enables scaling across multiple devices when required, making the implementation highly adaptable to various computational demands. The software accepts batches as input, with the batch size being user-defined and adjustable based on the sequence length and available GPU memory. Each batch of sequences is processed in parallel on GPU cores.

We employ `jax.lax.scan` as our primary loop construct instead of alternatives such as `jax.lax.fori loop` or `jax.lax.while loop`, due to its superior compatibility with just-in-time `jit` compilation. Unlike other constructs, `jax.lax.scan` is explicitly designed to facilitate efficient, parallelizable execution of iterative computations. By structuring the computation as a statically unrolled loop, `jax.lax.scan` not only minimizes overhead associated with dynamic control flow but also allows `JAX`’s `jit` compiler to optimize the entire computation graph in a single pass. This approach ensures that memory usage is reduced by efficiently handling intermediate states, a critical advantage when working with large datasets or long simulation runs. Furthermore, `jax.lax.scan` inherently supports reverse-mode automatic differentiation over iterative processes, making it particularly well-suited for gradient-based optimization tasks.

Figure 1 illustrates the schematic of our acceleration approach for Langevin-EPBD. All post-processing and data collection were performed on the CPU.

**Figure 1:**
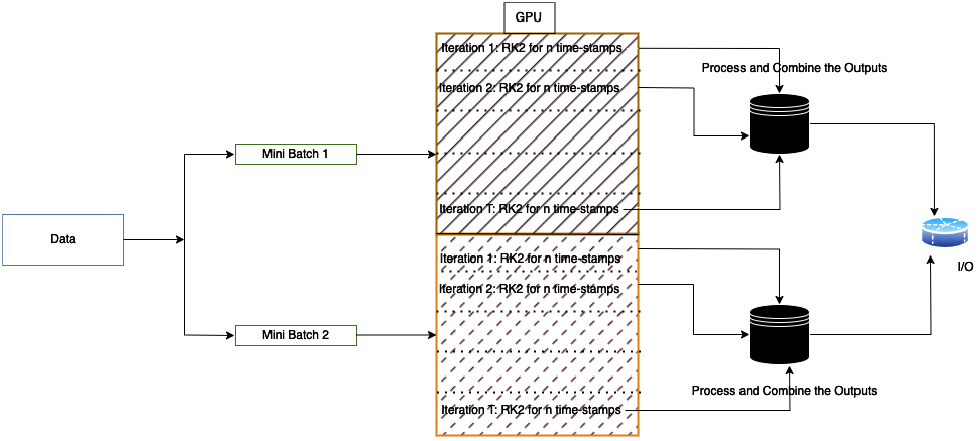
Schematic representation of the acceleration approach for *JAX-EPBD* using GPU parallelism. Multiple sequences are processed concurrently in batches on the GPU, while post-processing and data collection are performed on the CPU.

**Figure 2:**
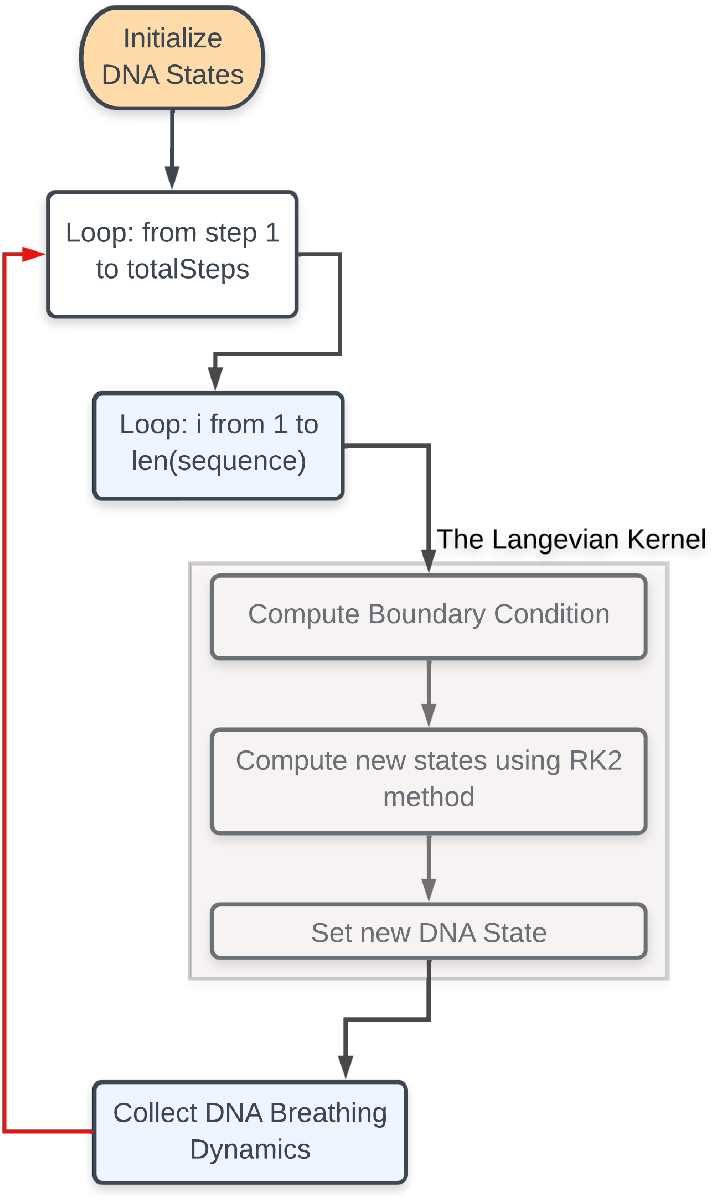
Langevin Dynamics Workflow in *JAX-EPBD* Framework

## Results and Discussion

First, we present the details of the datasets and simulation configurations used throughout the study. Next, we explore various aspects of DNA breathing dynamics, including base pair coordinates, base flipping, bubble formation, and the q-factor. We also highlight the utility of these breathing characteristics in predicting the binding specificity of Transcription Factors (TF) to DNA. Finally, we analyze the runtime performance of our JAX-EPBD model as a function of the number of base pairs in each DNA sequence.

### DataSets

We use several datasets to run the simulations and evaluate the DNA breathing dynamics features. This section discusses the datasets used in the study, which are essential for understanding the various utilities and perspectives of the Langevin-EPBD model.

#### Adeno-associated virus (AAV) P5 promoter

Experimental evidence suggests that spontaneous double-strand

DNA (dsDNA) separation at the transcriptional start site is a critical requirement for transcription initiation in several promoters (Alexandrov et al. 2009). This phenomenon, often referred to as DNA “breathing” or “bubble” formation, plays a pivotal role in creating an open complex that allows transcription machinery to bind and initiate RNA synthesis.

To investigate this process using our Langevin-EPBD model, we focus on the strand separation dynamics of the 77-base-pair-long AAV P5 promoter. This sequence serves as a key example of a promoter region where bubble dynamics are thought to play an essential role in transcriptional regulation. In addition to the wild-type promoter sequence, we study a control non-promoter sequence of the same length (77 bp). This mutant-type (mt) sequence contains a single mutation that alters the sequence’s ability to form bubbles, and it is derived from the published human collagen intron sequence (NW 927317). In figure 3 we present both the variants: wt and mt of P5 promoter; the mutation position is also highlighted.

**Figure 3:**
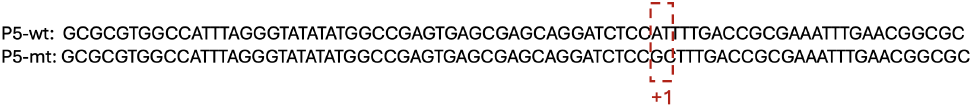
The sequence of Adeno-associated virus (AAV) P5 promoter. The mutation region is marked as +1 in the both wt and mt type)

#### SELEX

We utilize ground truth binding affinities obtained from high-throughput SELEX (HT-SELEX) experiments, focusing on specificity information for 215 transcription factors (TFs) from 27 families. The dataset, preprocessed by Yang et al. (Yang et al. 2017), includes TF-DNA binding specificities for all DNA sequences of length M (M-words) and underwent several filtering steps to ensure high variability, deep read coverage, selection of core motifs, and exclusion of infrequent M-words. This resulted in a comprehensive set of 1,788,827 sequences with lengths varying from 9 to 15 nucleotide base pairs. The affinity distribution shows similar variance across different TFs but shares the same maximum value, posing a significant challenge for computational learning models.

### Langevin-EPBD Simulation

We use the *JAX-EPBD* framework (see Figure 2) to monitor DNA breathing dynamics for a given DNA sequence. To ensure statistical significance, we perform at least 1,000 independent simulations with varying initial conditions (random seeds) at a constant temperature of *310 K*.

Each simulation consists of:

1. **Pre-heating Steps**: A preheating period of `200 picoseconds (ps)` allows the system to stabilize and remove initial condition artifacts.
2. **Simulation Phase**: We record dynamics for `1 nanosecond (ns)` with a time step of `1 femtosecond (fs)`. These durations are chosen based on convergence tests to adequately sample DNA breathing events. Other simulation parameters follow Alexandrov et al. (Alexandrov et al. 2006).

During simulations, base pair positions and velocities are updated using Langevin equations of motion, incorporating both deterministic forces from the EPBD potential and stochastic thermal fluctuations (Allen and Tildesley 1989). The integration time step is set to `0.00002` to balance accuracy and computational efficiency. We track the displacement *y*_*i*_ of each base pair *i* from its equilibrium position, indicating hydrogen bond stretching and DNA bubble formation. By averaging over multiple simulations, we obtain the average displacement profile ⟨*y*_*i*⟩_, which is sensitive to single base pair substitutions and does not require window averaging. For short DNA sequences, adding flanking regions is crucial to reflect the native base pair context, mitigate boundary effects from terminal base pairs, and satisfy the model’s minimum sequence length requirements. Our Langevin dynamics approach within the *JAX-EPBD* framework efficiently captures detailed, sequence-specific DNA breathing dynamics. By averaging numerous trajectories, we achieve precise profiles that enhance our understanding of the relationship between DNA structure, dynamics, and biological function. We will describe the DNA breathing dynamics below:

#### Breathing dynamics: Average coordinate distance profile

Average coordinates refer to the averaged transverse displacements, ⟨*y*_*i*_⟩, which are the displacements *y*_*i*_ averaged over thermal fluctuations. The displacement profile ⟨*y*_*i*_⟩ is a distinct characteristic of DNA breathing dynamics, quantifying the extent to which each base pair in the DNA sequence is “open” in equilibrium. This indicates the degree to which hydrogen bonds between base pairs are stretched due to thermal fluctuations.

Alexandrov et al. (Alexandrov et al. 2009) revealed that the simulation distribution correlates with significant differences in the transcriptional activity of promoters. To validate our approach, we analyzed the AAV P5 wild-type and mutant promoter sequences using the *JAX-EPBD* model. Figure 5f (right panel) displays the average displacements around base pair 50 for the wild-type promoter (blue) and a transcriptionally silent AG to TC mutant (red). These results are visually consistent with those reported by Alexandrov et al. (Alexandrov et al. 2009) and Kabir et al. (Kabir et al. 2023) (see figure 4). The average displacement magnitude in the double helix width can influence the binding affinity of transcription factors.

**Figure 4:**
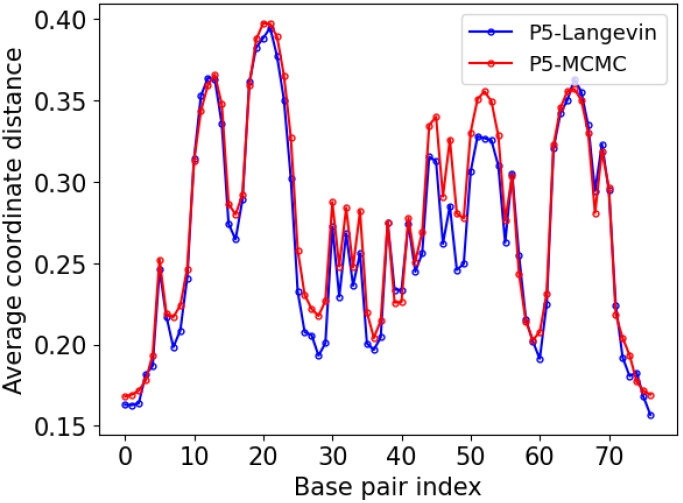
We visually show that average coordinate distance profile obtained by Langevin-EPBD is similiar to that of pyDNA-EPBD (Kabir et al. 2023)

**Figure 5:**
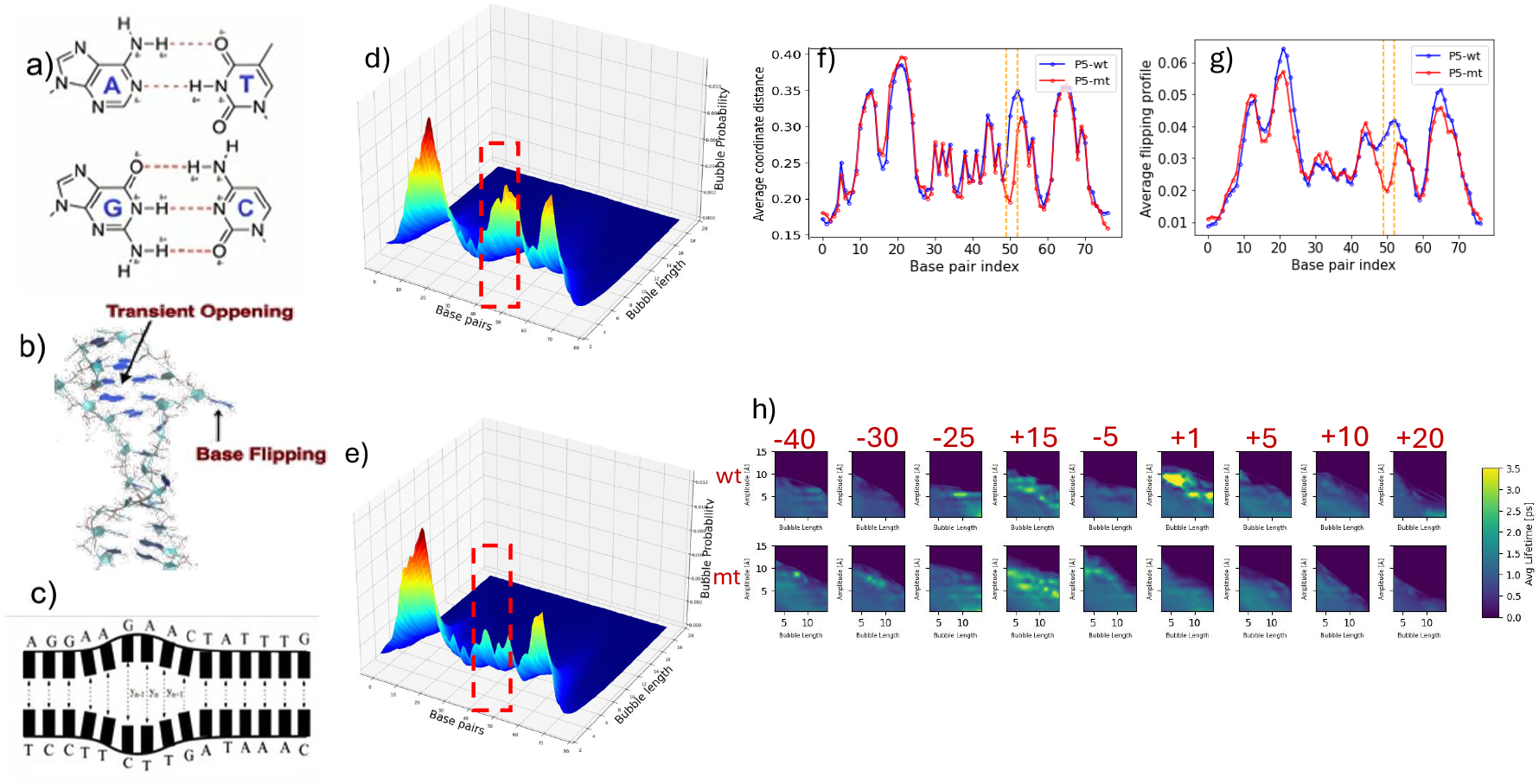
DNA Breathing Dynamics and Analysis. (a) The primary governance of macromolecules is through hydrogen bonds (H-bonds). (b) Representation of a single base “flipping out of the stack,” showcasing a phenomenon known as DNA breathing. (c) Illustration of consecutive base pairs breaking the H-bonds and opening simultaneously referred to as DNA bubbles. (d-e) 3D surface plots highlighting the change in bubble intensity across varied lengths and base pairs (bps) for threshold value 1.5 under two conditions: P5 wild (d) and P5 mutant (e). (f-g) Average Coordinates profiles for AAV P5 wild (f) and mutant-promoter (g) sequences at individual base pairs, with the orange vertical block indicating nucleotide substitutions from AG to TC at the 50 and 51 positions (zero-indexed). (h) Bubble lifetime for certain threshold as a function of bubble-length. The numbers in red indicates the distance from the mutation site in bp. +1 is the mutation site. For all the experiments, we set a minimum of 1000 Langevin simulations using various initial conditions, setting the temperature to 310 Kelvin and employing 200ps preheating steps followed by 1s post-preheating with 1fs time-step.

#### Breathing dynamics: Base Flipping Probability

Base flipping refers to a specific type of base movement in DNA where one or both bases in a base pair flip out of the helical stack, exposing them to the surrounding environment. This process is crucial for various biological functions, including DNA repair, replication, and transcription factor (TF) binding (Nowak-Lovato et al. 2013). The propensity for flipping characterizes this transition by determining the fraction of disrupted hydrogen bonds (openings) between complementary nucleotides. Specifically, it quantifies the fraction of base pairs (*bps*) whose displacement exceeds a certain threshold distance, as a function of temperature.

We computed the average flipping profile using our *JAX-EPBD* model by calculating the probability of a base pair being flipped throughout the simulation steps. A base pair is considered flipped if the separation between its bases equals or exceeds a predefined threshold distance (measured in A°). To accurately capture the flipping behavior, we collected flipping profiles at five different thresholds ranging from 0.7071, A° to 3.5355, A°, in increments of 0.7071, A° . Maintaining high floating-point precision is important to obtain accurate distribution profiles.

Figure 5g presents example flipping profiles for the wild-type and mutant adeno-associated virus (AAV) P5 promoter sequences at a threshold of 1.4142A° . The results show that the transcriptionally silent AAV P5 mutant is less prone to base pair openings at and around the mutation position at this threshold compared with its wild-type counterpart. This reduced propensity for flipping may contribute to the mutant’s transcriptional inactivity.

#### Breathing dynamics: Bubbles

DNA bubble probability refers to regions within the DNA double helix where the strands temporarily separate due to thermal motion. These transient denaturation bubbles are crucial for processes such as transcription initiation, replication, and transcription factor (TF) binding (Alexandrov et al. 2009; Choi et al. 2004). The formation of these bubbles is a stochastic process, especially in the presence of a thermal bath modeled by random forces (Nowak-Lovato et al. 2013). We define the probability of a DNA bubble, *P*_*n*_(*l, t*_*r*_), based on its starting base pair index *n*, length *l* (in base pairs), and displacement threshold *t*_*r*_ (in). This probability is expressed as:

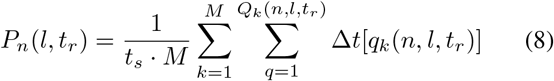

where:

1. *M* is the total number of simulation runs (typically *M≈* 1000).
2. *t*_*s*_ is the duration of a single simulation run, approximately 1 − 2 nanoseconds (ns).
3. *Q*_*k*_(*n, l, t*_*r*_) is the number of bubbles in the *k*-th simulation starting at base pair *n*, spanning *l* base pairs, and exceeding displacement *t*_*r*_.
4. Δ*t*[*q*_*k*_(*n, l, t*_*r*_)] is the existence time of the *q*-th bubble.

In the presence of the thermal bath, modeled by random forces, the creation of a bubble is a stochastic process (**?**). Unlike traditional MCMC methods, Langevin dynamics allow for time-dependent analyses, enabling the computation of bubble lifetimes. We compute average bubble life time as *τ*_Lifetime_, as:

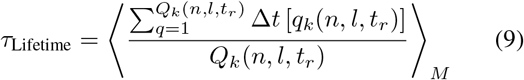

Using our *Langevin-EPBD* simulation tool, we analyzed DNA bubbles with lengths ranging from three to twenty base pairs and displacement thresholds from 0.5, A° to 15.0, A° in increments of 0.5, A° . High floating-point precision was maintained to ensure accurate distribution profiles.

In Figures 5d and 5e, we plot the bubble probability for a given threshold. Our results confirm previous findings (Choi et al. 2004; Alexandrov et al. 2006, 2009) that there is a significant difference in bubble probability at the mutation site between the two sequences, which corresponds to the dramatic difference in the transcriptional activity of the promoters. In Figure 5h, we present the average bubble lifetime for bubbles exceeding a certain amplitude as a function of bubble length. Consistent with earlier reports (Choi et al. 2004; Alexandrov et al. 2009), the P5 promoter displays a longer lifespan of bubbles at the mutation region.

#### Benchmark Results

In this section, we benchmark the performance of Jax-EPBD against the original C-based implementation (C-EPBD). All experiments were conducted on a computational node comprising 128 processors and an A100 GPU. The benchmarking experiments include two key scenarios: (1) varying sequence lengths for a fixed batch size, and (2) varying batch sizes for a fixed sequence length. These experiments aim to evaluate both runtime efficiency and scalability across different configurations. To establish a baseline comparison, we first evaluated the runtime for processing a single random sequence of length 100 across 100 simulations. The results are summarized in Table 1. Jax-EPBD achieved a significantly lower mean runtime of 7.21 seconds, compared to 67.80 seconds for C-EPBD, demonstrating approximately 9.4x speedup. Furthermore, Jax-EPBD exhibited greater consistency with a standard deviation of only 0.0135 seconds, compared to 0.1315 seconds for C-EPBD.

**Table 1:** Mean and standard deviation of runtime for Jax-EPBD and C-EPBD.

| Model | Mean | Standard Deviation |
| --- | --- | --- |
| Jax-EPBD | 7.2092 | 0.0135 |
| C-EPBD | 67.8000 | 0.1315 |

Next, we analyzed the runtime for a fixed batch size of 1 across varying sequence lengths, as illustrated in Figure 6. Jax-EPBD consistently demonstrated lower runtimes compared to C-EPBD, even as sequence lengths increased. Notably, the speedup metric—defined as the ratio of C-EPBD runtime to Jax-EPBD runtime—highlighted a growing performance advantage of Jax-EPBD for longer sequences.

**Figure 6:**
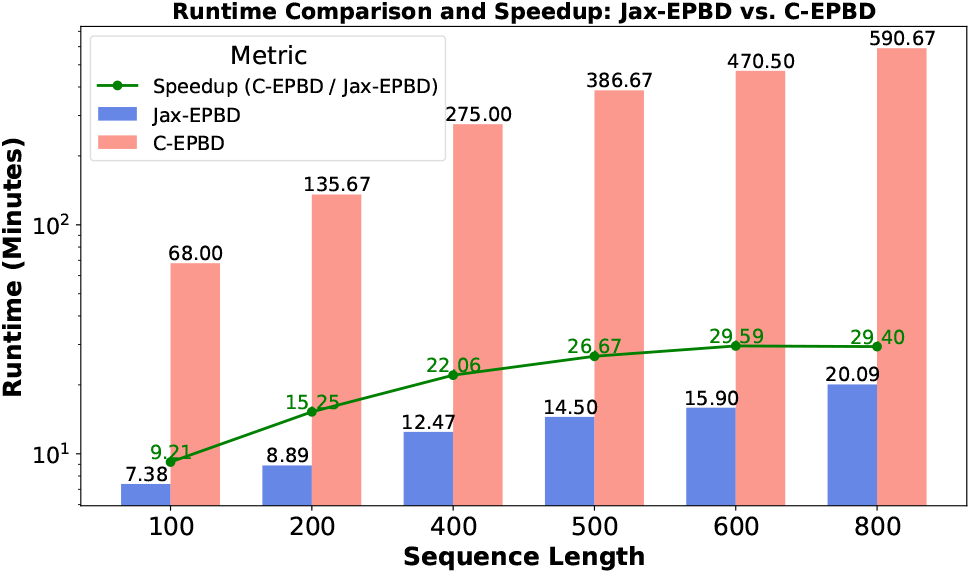
Runtime comparison and speedup analysis for Jax-EPBD and C-EPBD with a batch size of 1 across varying sequence lengths.

To evaluate the impact of increasing batch size, we compared the runtime for a fixed sequence length of 100 across various batch sizes, as presented in Figure 7. Jax-EPBD exhibited significantly lower runtimes than C-EPBD, with consistent scaling as batch size increased. In contrast, C-EPBD showed progressively higher runtimes and became infeasible for batch sizes beyond 40 due to resource limitations. The speedup metric further emphasized the efficiency gains of Jax-EPBD, particularly at larger batch sizes, where its performance remained stable. For example, at a batch size of 40, Jax-EPBD achieved a runtime of 110.05 seconds, compared to 2025.60 seconds for C-EPBD. These findings highlight the adaptability and computational efficiency of Jax-EPBD in scenarios involving high-throughput simulations making it particularly well-suited for large-scale genomic applications.

**Figure 7:**
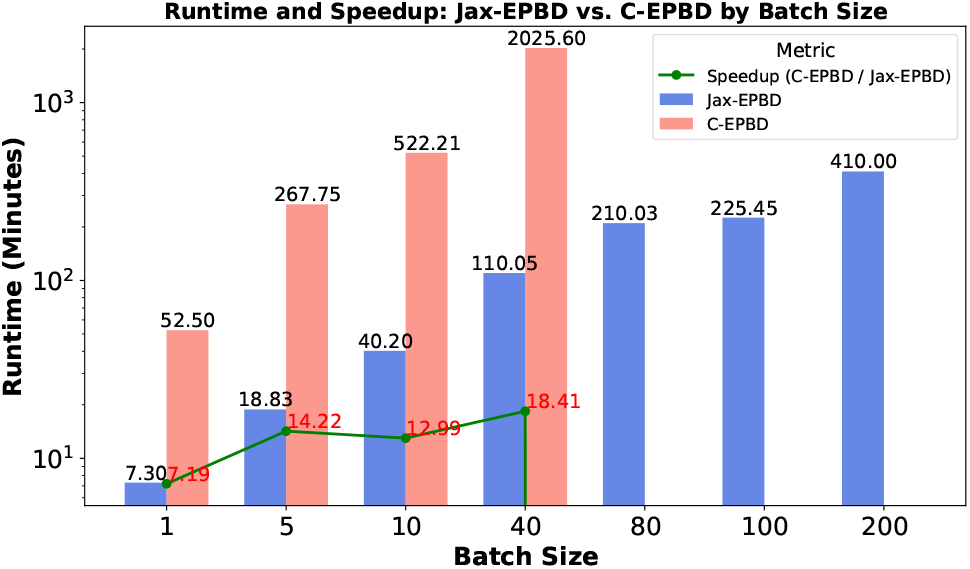
Runtime and speedup comparison between Jax-EPBD and C-EPBD for sequence length 100 across various batch sizes.

#### Binding prediction

To analyze transcription factor (TF) binding affinity predictions on the SELEX dataset, we leveraged a simple Support Vector Regression (SVR) model with a linear kernel and evaluated its performance using 10-fold cross-validation. Initially, the model was evaluated using only the one-hot encoded representation of the DNA sequences. Subsequently, we extended the feature set to include sequence information combined with DNA breathing features, primarily coordinate features derived from structural dynamics. Our key observations were as follows: the inclusion of breathing features consistently improved predictive performance across most TF families, as indicated by higher *R*^2^ values as shown in Figure 8. For example, the “C2H2” zinc finger family and homeodomain family demonstrated significant gains in prediction accuracy with the addition of breathing features, highlighting their importance in capturing subtle sequence-dependent interactions. In contrast, TF families with inherently strong baseline performance from one-hot encoding, such as certain nuclear receptor proteins, exhibited relatively modest improvements.

**Figure 8:**
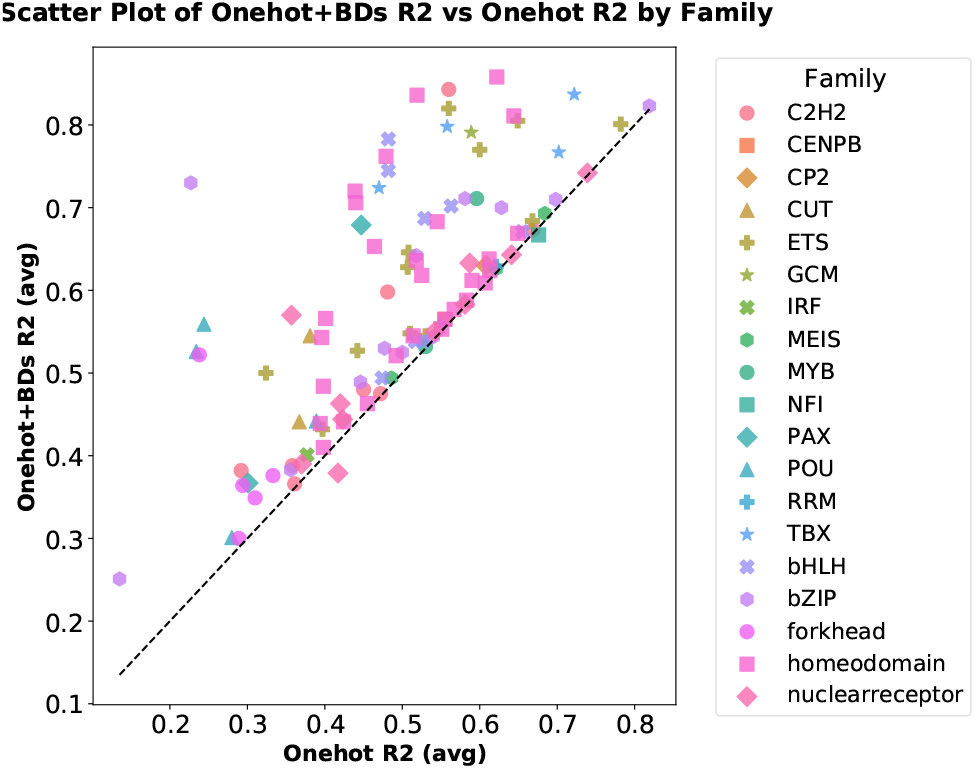
Scatter plot comparing Onehot *R*^2^ (average) and Onehot+BDs *R*^2^(average) across different Transcription factor families. Each family is represented by a unique combination of marker style and color, highlighting variations in performance. The dashed black line represents the ideal relationship (y = x), indicating parity between the two methods

## Conclusion

We developed and implemented *JAX-EPBD*, a high-throughput Langevin molecular dynamics (LMD) simulation framework for studying DNA breathing dynamics with unprecedented efficiency and detail. Leveraging the computational power of the `JAX` library and GPU acceleration, our framework overcomes the temporal resolution and scalability limitations of traditional methods like MCMC. *JAX-EPBD* achieves exceptional runtime efficiency, offering up to 30x speedup over the original C-based EPBD implementation, and scales effectively across varying sequence lengths and batch sizes. This scalability enables the processing of multiple DNA sequences concurrently, accommodating longer sequences and capturing rare breathing events critical for comprehensive DNA dynamics studies.

The framework’s capabilities were validated using the 77-base-pair AAV P5 promoter, where we detected subtle but significant differences in breathing dynamics between wild-type and mutant sequences. These differences, reflected in metrics such as average coordinate displacements, base flipping probabilities, and bubble lifetimes, aligned with experimental observations of transcriptional activity, further confirming the biological relevance of our approach. Additionally, the integration of DNA breathing features into transcription factor (TF) binding affinity predictions demonstrated the utility of *JAX-EPBD* beyond fundamental dynamics, with enhanced *R*^2^ values across most TF families, including the “C2H2” zinc finger and homeodomain families. *JAX-EPBD* not only provides a robust and scalable platform for studying DNA breathing dynamics but also offers insights into how sequence variations influence genetic function. Its efficiency and scalability make it a valuable tool for exploring transcription factor binding specificity, DNA repair mechanisms, and the effects of single nucleotide polymorphisms on DNA stability. This advancement opens new avenues for genome-scale analyses and deeper investigations into the fundamental mechanisms governing genetic processes.

## Supporting information

Langevin Dynamics Workflow in JAX-EPBD Framework

## Acknowledgements

This research project was supported by the National Institute of Health (NIH) RO1HL128831 (to A.U.) and the NIH 5R01MH116281-03 (to B.A.). M.B. and T.I. were supported by NIH 5R01MH116281-03 and the LANL LDRD 20230287ER grant. A.K. and A.S. were supported in part by the National Science Foundation Grant No. 2310113. This research used resources provided by the Los Alamos National Laboratory Institutional Computing Program, which is supported by the U.S. Department of Energy National Nuclear Security Administration under Contract No. 89233218CNA000001.

## References

Alexandrov, B.; Wille, L.; Rasmussen, K.; Bishop, A.; and Blagoev, K. 2006. Bubble statistics and dynamics in double-stranded DNA. Physical Review E—Statistical, Nonlinear, and Soft Matter Physics, 74(5): 050901.

Alexandrov, B. S.; Gelev, V.; Bishop, A. R.; Usheva, A.; and Rasmussen, K.Ø. 2010a. DNA breathing dynamics in the presence of a terahertz field. Physics Letters A, 374(10): 1214–1217.

Alexandrov, B. S.; Gelev, V.; Monisova, Y.; Alexandrov, L. B.; Bishop, A. R.; Rasmussen, K.Ø.; and Usheva, A. 2009. A nonlinear dynamic model of DNA with a sequence-dependent stacking term. Nucleic Acids Research, 37(7): 2405–2410.

Alexandrov, B. S.; Gelev, V.; Yoo, S. W.; Alexandrov, L. B.; Fukuyo, Y.; Bishop, A. R.; Rasmussen, K.Ø.; and Usheva, A. 2010b. DNA dynamics play a role as a basal transcription factor in the positioning and regulation of gene transcription initiation. Nucleic acids research, 38(6): 1790–1795.

Allen, M. P.; and Tildesley, D. J. 1989. Computer Simulation of Liquids. Oxford: Oxford University Press. ISBN 978-0198556459.

Ares, S.; Voulgarakis, N.; Rasmussen, K.; and Bishop, A. R. 2005. Bubble nucleation and cooperativity in DNA melting. Physical review letters, 94(3): 035504.

Bradbury, J.; Frostig, R.; Hawkins, P.; Johnson, M. J.; Leary, C.; Maclaurin, D.; Necula, G.; Paszke, A.; VanderPlas, J.; Wanderman-Milne, S.; and Zhang, Q. 2018. JAX: composable transformations of Python+NumPy programs.

Choi, C. H.; Kalosakas, G.; Rasmussen, K.Ø.; Hiromura, M.; Bishop, A. R.; and Usheva, A. 2004. DNA dynamically directs its own transcription initiation. Nucleic Acids Research, 32(4): 1584–1590.

Frenkel, D.; and Smit, B. 2002. Understanding Molecular Simulation: From Algorithms to Applications. San Diego: Academic Press, 2nd edition. ISBN 978-0122673511.

Guèron, M.; Kochoyan, M.; and Leroy, J.-L. 1987. A single mode of DNA base-pair opening drives imino proton exchange. Nature, 328(6125): 89–92.

Ixaru, L. G.; Vanden Berghe, G.; Ixaru, L. G.; and Vanden Berghe, G. 2004. Runge-Kutta Solvers for Ordinary Differential Equations. Exponential Fitting, 223–304.

Kabir, A.; Bhattarai, M.; Rasmussen, K. ; Shehu, A.; Usheva, A.; Bishop, A. R.; and Alexandrov, B. 2023. Examining DNA breathing with pyDNA-EPBD. Bioinformatics, 39(11): btad699.

Nowak-Lovato, K.; Alexandrov, L. B.; Banisadr, A.; Bauer, A. L.; Bishop, A. R.; Usheva, A.; Mu, F.; Hong-Geller, E.; Rasmussen, K.Ø.; Hlavacek, W. S.; et al. 2013. Binding of nucleoid-associated protein Fis to DNA is regulated by DNA breathing dynamics. PLoS Computational Biology, 9(1): e1002881.

Peyrard, M. 2004. Nonlinear dynamics and statistical physics of DNA. Nonlinearity, 17(2): R1.

Peyrard, M.; and Bishop, A. R. 1989. Statistical mechanics of a nonlinear model for DNA denaturation. Physical review letters, 62(23): 2755.

SantaLucia Jr, J. 1998. A unified view of polymer, dumbbell, and oligonucleotide DNA nearest-neighbor thermodynamics. Proceedings of the National Academy of Sciences, 95(4): 1460–1465.

Watson, J. D.; Baker, T. A.; Bell, S. P.; Gann, A.; Levine, M.; and Losick, R. 2013. Molecular Biology of the Gene. New York: Pearson, 7th edition. ISBN 978-0321762436.

Yang, L.; Orenstein, Y.; Jolma, A.; Yin, Y.; Taipale, J.; Shamir, R.; and Rohs, R. 2017. Transcription factor family-specific DNA shape readout revealed by quantitative specificity models. Molecular systems biology, 13(2): 910.

