## Supplementary figures and images for "Efficient High-Throughput DNA Breathing Features Generation Using Jax-EPBD"

### Langevin Dynamics Workflow in JAX-EPBD Framework

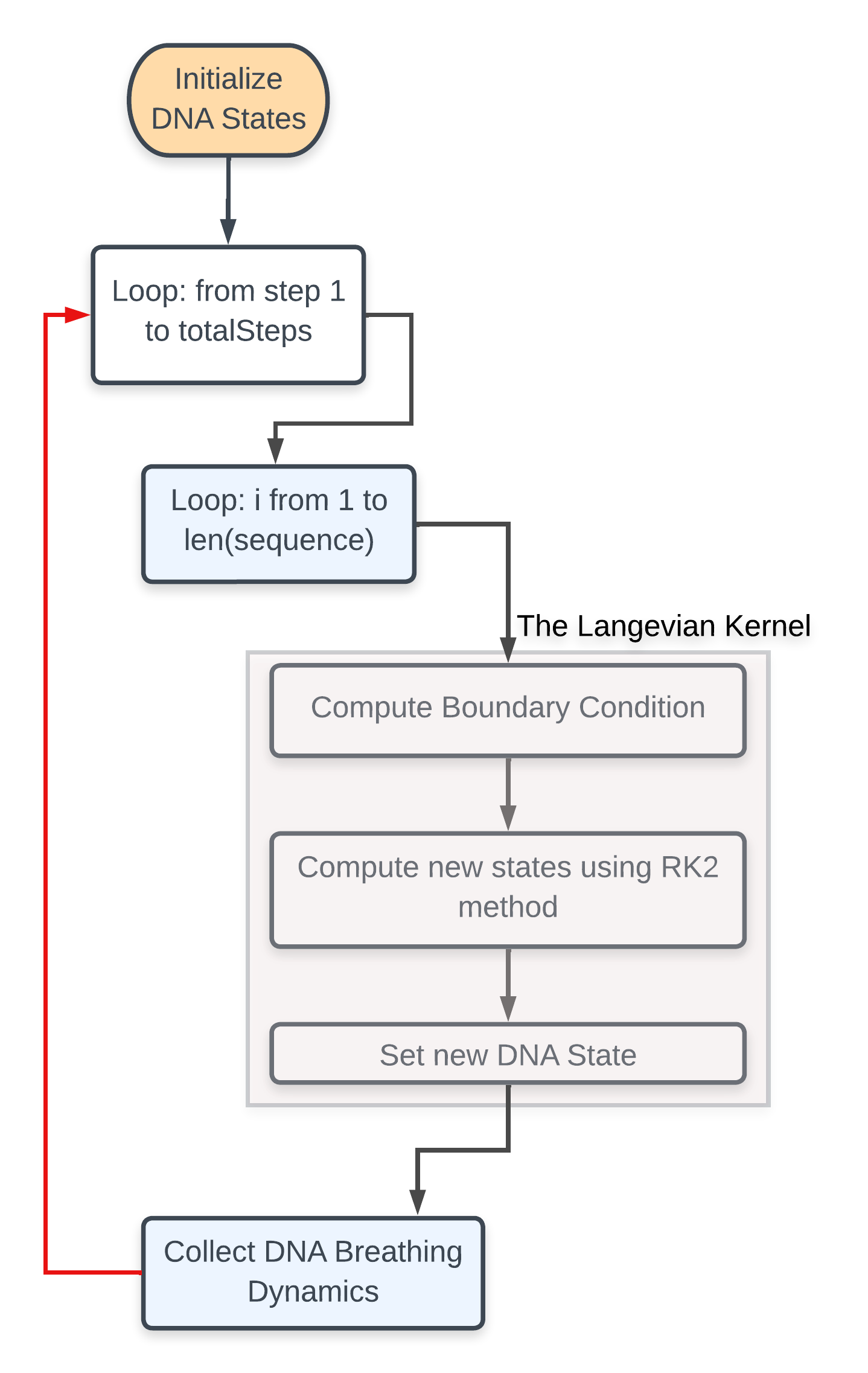
